# TMS disruption of the lateral prefrontal cortex increases neural activity in the default mode network when naming facial expressions

**DOI:** 10.1101/2023.03.09.531897

**Authors:** David Pitcher, Magdalena W. Sliwinska, Daniel Kaiser

## Abstract

Recognizing facial expressions is dependent on multiple brain networks specialized for different cognitive functions. In the current study participants (*N*=20) were scanned using functional magnetic resonance imaging (fMRI) while they performed a covert facial expression naming task. Immediately prior to scanning thetaburst transcranial magnetic stimulation (TMS) was delivered over the right lateral prefrontal cortex (PFC), or the vertex control site. A group whole-brain analysis revealed that TMS induced opposite effects in the neural responses across different brain networks. Stimulation of the right PFC (compared to stimulation of the vertex) decreased neural activity in the left lateral PFC but increased neural activity in three nodes of the default mode network (DMN): the right superior frontal gyrus (SFG), right angular gyrus and the bilateral middle cingulate gyrus. A region of interest (ROI) analysis showed that TMS delivered over the right PFC reduced neural activity across all functionally localised face areas (including in the PFC) compared to TMS delivered over the vertex. These results causally demonstrate that visually recognizing facial expressions is dependent on the dynamic interaction of the face processing network and the DMN. Our study also demonstrates the utility of combined TMS / fMRI studies for revealing the dynamic interactions between different functional brain networks.

## Introduction

Humans need to recognize and interpret the facial expressions of other people during social interactions. The neural computations that support these cognitive processes have been extensively investigated using functional magnetic resonance imaging (fMRI). These studies have been the basis of theories positing that emotions are processed across multiple large scale brain networks that engage both cortical and subcortical structures (Barrett & Satpute, 2013; Lindquist et al., 2012; Pessoa, 2018; Wager et al., 2015). The extent to which these networks interact with brain networks specialized for other cognitive functions has also been investigated. For example, it has been proposed that emotion processing is reliant on dynamic interactions between the salience network (e.g., the amygdala and insula) and the central executive brain network for cognitive control (e.g., the lateral prefrontal cortex) (Pessoa, 2018; Seeley et al., 2007; Uddin, 2015). Both the salience network and the central executive network consist of brain areas that show greater neural activation when participants perform tasks requiring emotional processing. However, a recent model has proposed that another brain network, the default mode network (DMN) is also necessary for emotion processing (Satpute & Lindquist, 2019).

The DMN is anti-correlated with task performance, meaning it exhibits a decrease in neural activity when participants perform cognitive tasks in the fMRI scanner (Raichle, 2015). This has led to claims that the DMN mediates inner states such as mind wandering, inner thoughts and internal states (Smallwood et al., 2021). fMRI studies have also demonstrated that the DMN is anti-correlated with other brain areas during facial expression naming tasks (Lanzoni et al., 2020; Sreenivas et al., 2012). This is consistent with the hypothesis that emotion processing is dependent on a push / pull interaction between the salience and central executive networks and the DMN (Satpute & Lindquist, 2019). Our aim in the current study was to causally test the role of the DMN in a facial expression naming task by combining fMRI with transcranial magnetic stimulation (TMS). To do this we transiently disrupted the right lateral prefrontal cortex (PFC), a node in the central executive network that is anti-correlated with the DMN (Raichle, 2015).

The lateral PFC is involved in a range of different face processing tasks including identity recognition (Ishai et al., 2002), working memory for faces (Courtney et al., 1996) and the configural processing of the eyes and mouth (Renzi et al., 2013). Importantly, prior studies have also demonstrated that the lateral PFC is involved in facial expression processing (Gorno-Tempini et al., 2001; Iidaka et al., 2001). Neuropsychological studies of patients with frontal lobe damage have further demonstrated that those with lateral PFC damage have problems with a range of emotion processing tasks including theory of mind and self-emotion regulation (Jastorff et al., 2016; Tsuchida & Fellows, 2012). Based on these studies we chose to disrupt the PFC with TMS while participants performed a facial expression naming task in the fMRI scanner. TMS was delivered over the inferior frontal gyrus (IFG), a region of the lateral PFC that has been implicated in a range of face processing tasks (Chan, 2013; Ishai et al., 2002). We chose to target the right IFG because our prior study demonstrated that face-selective activity can be more reliably identified in the right hemisphere (Nikel et al., 2022).

Patient studies have also demonstrated that facial expression recognition is dependent on a wider network of visual brain areas that selectively process faces (Adolphs, 2002; Jastorff et al., 2016). These include areas of the temporal cortex that are known to contain face-selective areas in both the ventral (Kanwisher et al., 1997; McCarthy et al., 1997) and lateral (Gauthier et al., 2000; Puce et al., 1996; Puce et al., 1998) brain surfaces. These areas have been linked together into models that propose a distributed brain network specialized for face processing (Calder & Young, 2005; Haxby et al., 2000). Prior neuroimaging studies have also revealed that the lateral PFC is engaged in the top-down control of other brain areas when recognizing faces including the amygdala (Davies-Thompson & Andrews, 2012), ventral temporal cortex (Baldauf & Desimone, 2014; Heekeren et al., 2004) and the superior temporal cortex (STS) (Wang et al., 2020). Connectivity between face areas has also been causally demonstrated. Our prior combined TMS/fMRI studies revealed that disruption of one node in the face network impaired neural activity in remote face areas (Groen et al., 2021; Pitcher et al., 2014; Pitcher et al., 2017) and the functional connectivity between these areas (Handwerker et al., 2020). We therefore predicted that TMS delivered over the right PFC while participants performed a facial expression naming task would decrease neural activity across the face network. Crucially we also predicted that transient disruption of this network would cause an increase in neural activity in the DMN because the two networks dynamically interact during facial expression naming.

## Materials and Methods

### Participants

A total of 20 participants (14 females; age range 19 to 46 years old; mean age 23 years, SD = 6.4) with normal or corrected-to-normal vision gave informed consent as directed by the Ethics Committee at the University of York.

### Stimuli

Face stimuli for the expression naming task were 14 models (female and male) from Ekman and Friesen’s (1976) facial affect series expressing one of seven emotions. Each image was shown once only. This equated to a total of 110 unique pictures: anger (17), disgust (15), fear (17), happy (18), neutral (14), sad (15) and surprise (14).

In addition to the experimental task, we also ran a functional localizer to identify face-selective areas in each participant. Stimuli were 3-second movie clips of faces and objects that we have used in prior studies (Küçük et al., 2022; Sliwinska, Elson, et al., 2020; Sliwinska et al., 2022). There were sixty movie clips for each category in which distinct exemplars appeared multiple times. Movies of faces and bodies were filmed on a black background, and framed close-up to reveal only the faces or bodies of 7 children as they danced or played with toys or adults (who were out of frame). Fifteen different moving objects were selected that minimized any suggestion of animacy of the object itself or of a hidden actor pushing the object (these included mobiles, windup toys, toy planes and tractors, balls rolling down sloped inclines). Within each block, stimuli were randomly selected from within the entire set for that stimulus category. This meant that the same actor or object could appear within the same block but given the number of stimuli this did not occur regularly.

### Procedure

Participants completed three separate fMRI sessions, each performed on a different day. The first session was an fMRI experiment designed to individually localize the TMS sites in each participant. In this session participants viewed three runs of a functional localiser task (234 seconds each) to individually identify face-selective areas. Our previous fMRI study of face processing in the lateral PFC demonstrated that a face-selective area was more commonly identified across participants in the right IFG (Nikel et al., 2022). Based on this study we targeted the same location for disruption with TMS in the current study. Functional runs presented short video clips of faces, bodies and objects in 18-second blocks that contained six 3-second video clips from that category. We also collected a high-resolution structural scan for each participant. The remaining two sessions were combined TMS/fMRI sessions.

### Combined TMS/fMRI Sessions

Prior to the combined TMS/fMRI we used the Brainsight TMS-MRI co-registration system (Rogue Research) to mark the location of the face-selective area in the right IFG based on the initial fMRI localizer data collected for each participant. The vertex control site was identified using a tape measure as a point in the middle of the head halfway between the nasion and inion (Figure 1a).

**Figure 1a.**
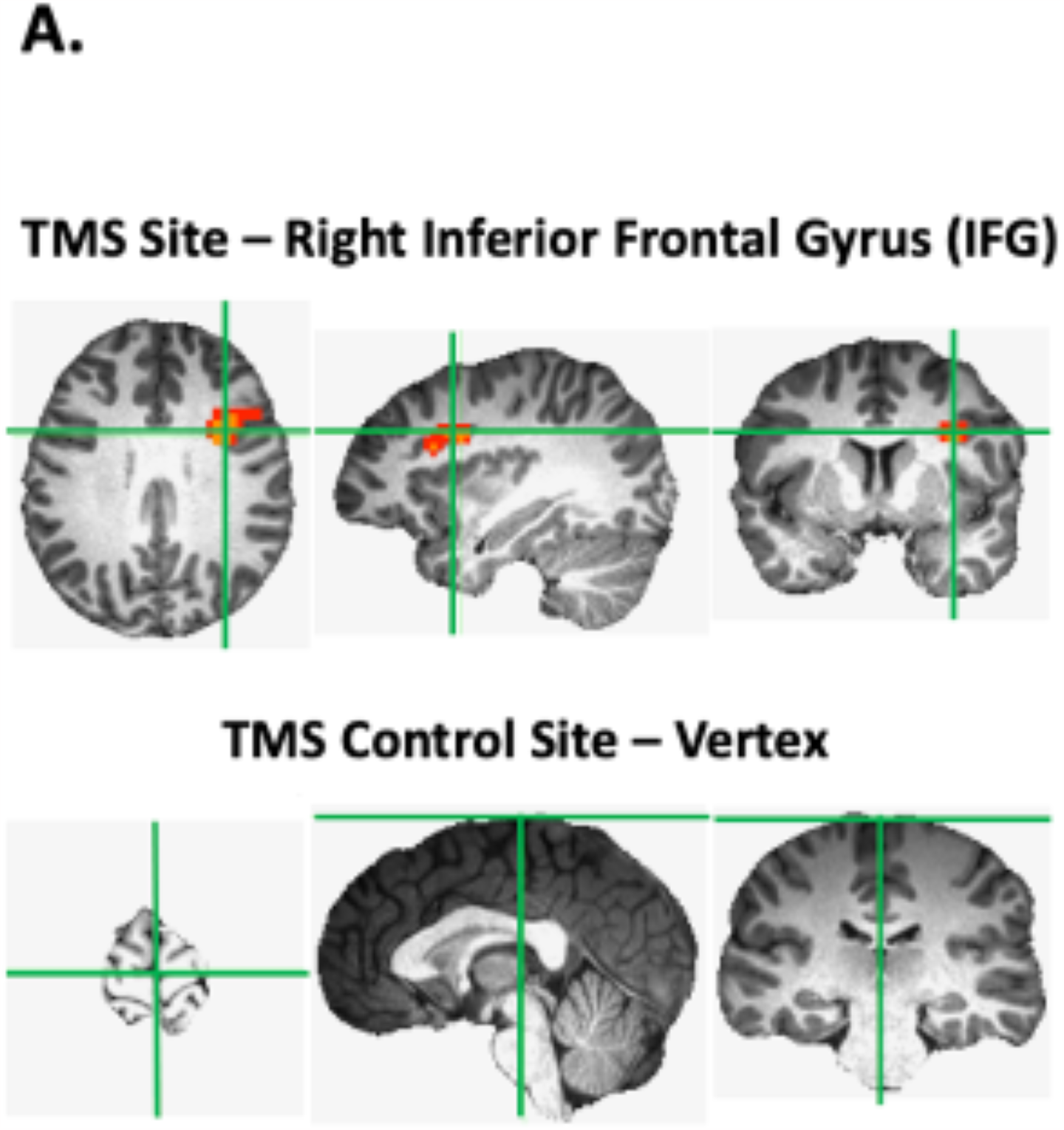
The TMS sites from an example participant. The active site in the right lateral PFC was defined using a contrast of faces > objects for each participant. The average MNI coordinates (36,6,48) were centred in right IFG.

Participants were then taken to the fMRI scanner control room where thetaburst TMS was delivered over the right IFG or the vertex for each participant (stimulation site order was balanced across participants). Once stimulation was completed, participants entered the scanner room immediately. fMRI data collection began as quickly as possible as the effects of TMS are transient. For all participants, scanning started within five minutes of TMS stimulation being delivered.

Functional data for the expression naming task were acquired over 2 blocked-design functional runs lasting 570 seconds each (Figure 1b). Each run consisted of 55 trials during which facial expression stimuli were presented centrally on the screen for 3 seconds, followed by a blank screen of 7 seconds. Participants were instructed to silently name the emotion that the facial expression displayed when the stimuli was presented. Once the two expression naming runs were completed participants viewed two runs of the localizer task (234 seconds each) to individually identify face-selective areas. Functional runs presented short video clips of faces, bodies and objects in 18-second blocks that contained six 3-second video clips from that category. Once localizer data collection was completed participants exited the scanner. After the final session participants were debriefed on the nature of the study.

### Brain Imaging and Analysis

Imaging data were acquired using a 3T Siemens Magnetom Prisma MRI scanner (Siemens Healthcare, Erlangen, Germany) at the University of York. Functional images were acquired with a twenty-channel phased array head coil and a gradient-echo EPI sequence (38 interleaved slices, repetition time (TR) = 3 sec, echo time (TE) = 30ms, flip angle =90 degrees; voxel size 3mm isotropic; matrix size = 128 × 128) providing whole brain coverage. Slices were aligned with the anterior to posterior commissure line. Structural images were acquired using the same head coil and a high-resolution T-1 weighted 3D fast spoilt gradient (SPGR) sequence (176 interleaved slices, repetition time (TR) = 7.8 sec, echo time (TE) = 3ms, flip angle = 20 degrees; voxel size 1mm isotropic; matrix size = 256 × 256).

Functional MRI data were analyzed using AFNI (http://afni.nimh.nih.gov/afni). Data from the first four TRs from each run were discarded. The remaining images were slice-time corrected and realigned to the last volume of the last run prior to TMS during the TMS to vertex session, and to the corresponding anatomical scan. The volume registered data were spatially smoothed with an 8mm full-width-half-maximum Gaussian kernel. Signal intensity was normalized to the mean signal value within each run and multiplied by 100 so that the data represented percent signal change from the mean signal value before analysis.

For the expression naming task, we performed two separate analyses. The first grouped all expressions together using a general linear model (GLM) was established by convolving the standard hemodynamic response function with one regressor of interest (faces). In the second analysis we examined each of the seven expressions separately using a general linear model (GLM) was established by convolving the standard hemodynamic response function with seven regressor of interest (angry, disgust, fear, happy, neutral, sad and surprise). For both analyses regressors of no interest (e.g., 6 head movement parameters obtained during volume registration and AFNI’s baseline estimates) were also included in each GLM.

For the localizer task a general linear model (GLM) was established by convolving the standard hemodynamic response function with two regressors of interest (faces and objects). Regressors of no interest (e.g., 6 head movement parameters obtained during volume registration and AFNI’s baseline estimates) were also included.

We also performed an exploratory multi-voxel pattern analysis (MVPA) to determine whether TMS delivered over the right IFG decreases the neural discriminability between different expressions in the face network. This analysis was performed for IFG, the Amygdala, pSTS, FFA, and OFA. Here, we used ROI masks that included both hemispheres to increase signal-to-noise ratio in the light of the limited data available. We first created new GLMs, which contained seven regressors for each of the seven emotions (anger, disgust, fear happy, neutral, sad and surprise), separately for each fMRI run. From these GLMs, we then calculated T-maps against baseline for each emotion. The subsequent MVPA analysis was carried out using the CoSMoMVPA toolbox for Matlab (Oosterhof et al., 2016). To quantify the discriminability between emotions, we used a cross-validated correlation approach (Haxby et al., 2001). Specifically, we correlated (*Spearman*-correlation) response patterns (T-values across voxels in each ROI) between the two runs, either for the same emotion (within-correlations) or for different emotions (between-correlations). Subtracting the between-correlations from the within-correlations yielded a measure of neural discriminability between emotions for each ROI, separately for the two TMS sites.

### TMS Site Localization and parameters

Stimulation sites were localized using individual structural and functional images collected during an fMRI localizer task that each participant completed prior to the combined TMS/fMRI sessions. In the localizer session, participants viewed the same dynamic face and object stimuli as in earlier studies of the face network (Pitcher et al., 2011; Sliwinska, Bearpark, et al., 2020). The stimulation site targeted in the right IFG (Nikel et al., 2022) of each participant was the peak voxel in the face-selective ROI identified using a contrast of greater activation by dynamic faces than dynamic objects (mean MNI co-ordinates 36,6,48). The vertex site was identified as a point on the top of the head halfway between the nasion (the tip of the nose) and the inion (the point at the back of the head). TMS sites were identified using the Brainsight TMS-MRI co-registration system (Rogue Research) and the proper coil locations were then marked on each participant’s scalp using a marker pen.

A Magstim Super Rapid Stimulator (Magstim; Whitland, UK) was used to deliver the TMS via a figure-eight coil with a wing diameter of 70 mm. TMS was delivered at an intensity of 45% of machine output over each participant’s functionally localized right IFG or vertex. Thetaburst TMS (TBS) was delivered using a continuous train of 600 pulses delivered in bursts of 3 pulses (a total of 200 bursts) at a frequency of 30 Hz with a burst frequency of 6 Hz for a duration of 33.3 sec and fixed intensity of 45% of the maximum stimulator output. We used a modified version (Nyffeler et al., 2006) of the original thetaburst protocol (Huang et al., 2005) as this version has been shown to have longer lasting effects (Goldsworthy et al., 2012). The Stimulator coil handle was held pointing upwards and parallel to the midline when delivered over the right IFG and flat against the skull with handle towards the inion when delivered over the vertex.

### Data Availability Statement

The data underlying this article will be shared on reasonable request to the corresponding author.

## Results

### Whole brain group analysis of TMS disruption of the right IFG

Experimental data (*N*=20) from the expression naming task were entered into a group whole brain ANOVA. Activation maps were calculated for each TMS session and the IFG session data were then subtracted from the vertex session data (*p*=0.005, zstat = 3.1). These maps were then registered to the MNI template using probabilistic maps for combining functional imaging data with cytoarchitectonic maps (Eickhoff et al., 2005). Results revealed multiple brain areas that exhibited significant differences between TMS sites (Figure 2a). TMS delivered over the right IFG reduced neural activity in the left IFG (-46, 22, 17) while increasing neural activity in three nodes of the default mode network: the right superior frontal gyrus (SFG) (20, 37, 35), right angular gyrus (47, -50, 29) and bilateral middle cingulate cortex (5, -23, 38). To determine response magnitudes against baseline, we also calculated the percent signal change for the two stimulation conditions in these four regions (Figure 2b). Consistent with the established neural response pattern of the DMN we observed a negative BOLD response in the right SFG, right angular gyrus and bilateral cingulate cortex in the vertex control condition. However, when we disrupted the salience network after TMS was delivered over the right IFG, the negativity of the BOLD response was reduced in all three areas. By contrast, TMS delivered over the right IFG revealed the opposite effect on the neural activity in the left IFG. Namely, disruption of the right IFG reduced the positive neural activity in the left IFG when performing the facial expression naming task (Figure 2b).

**Figure 2a.**
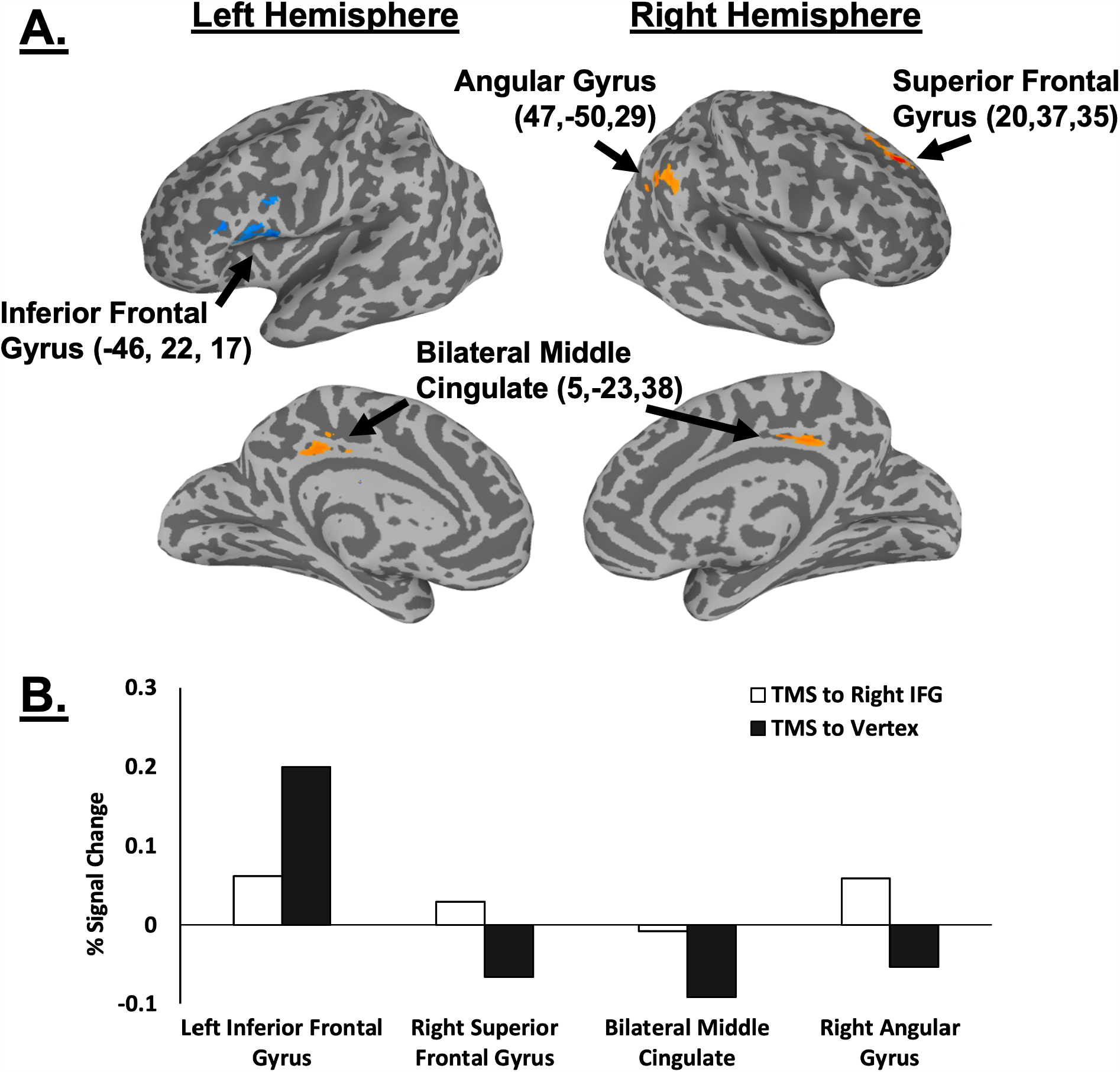
The results of a group whole brain analysis showing the distributed impact of TMS delivered over the right IFG while participants silently named facial expressions. Group data (*N*=20) where calculated for each TMS session and the IFG session data were then subtracted from the vertex session data (*p*=0.005, zstat = 3.1). Clusters in orange denote an increase in neural activity after TMS delivered over the right IFG. The cluster in blue denote a decrease in neural activity after TMS delivered over the right IFG. Figure 2b. The percent signal change for the two stimulation conditions in the four regions identified in the group analysis. TMS delivered over the right IFG reduced the positive neural activity in the left IFG and increased the negative neural activity in the right SFG, right angular gyrus and bilateral middle cingulate cortex (components of the default mode network).

### Region of Interest (ROI) analysis of TMS disruption in face processing network

To further characterise the effects of disrupting the right IFG across the face processing network we also performed a region of interest (ROI) analysis at the individual participant level. ROIs were defined using the functional localiser runs from the initial fMRI session and the runs collected after the expression naming task in the combined TMS/fMRI sessions. Face-selective ROIs were identified across both hemispheres using a contrast of faces greater than objects and a statistical threshold of *p* < 0.1. This was based on our prior study of the face-selective areas in the bilateral IFG which demonstrated that this threshold was necessary (Nikel et al., 2022). We identified clusters of at last 5 voxels in each defined face area and created a 5mm sphere around the peak activation coordinate for the following ROIs in both hemispheres: IFG, Amygdala, posterior superior temporal sulcus (pSTS), fusiform face area (FFA) and occipital face area (OFA). We then calculated the percent signal change for the two stimulation conditions (right IFG and vertex) in each ROI (Figure 3).

**Figure 3.**
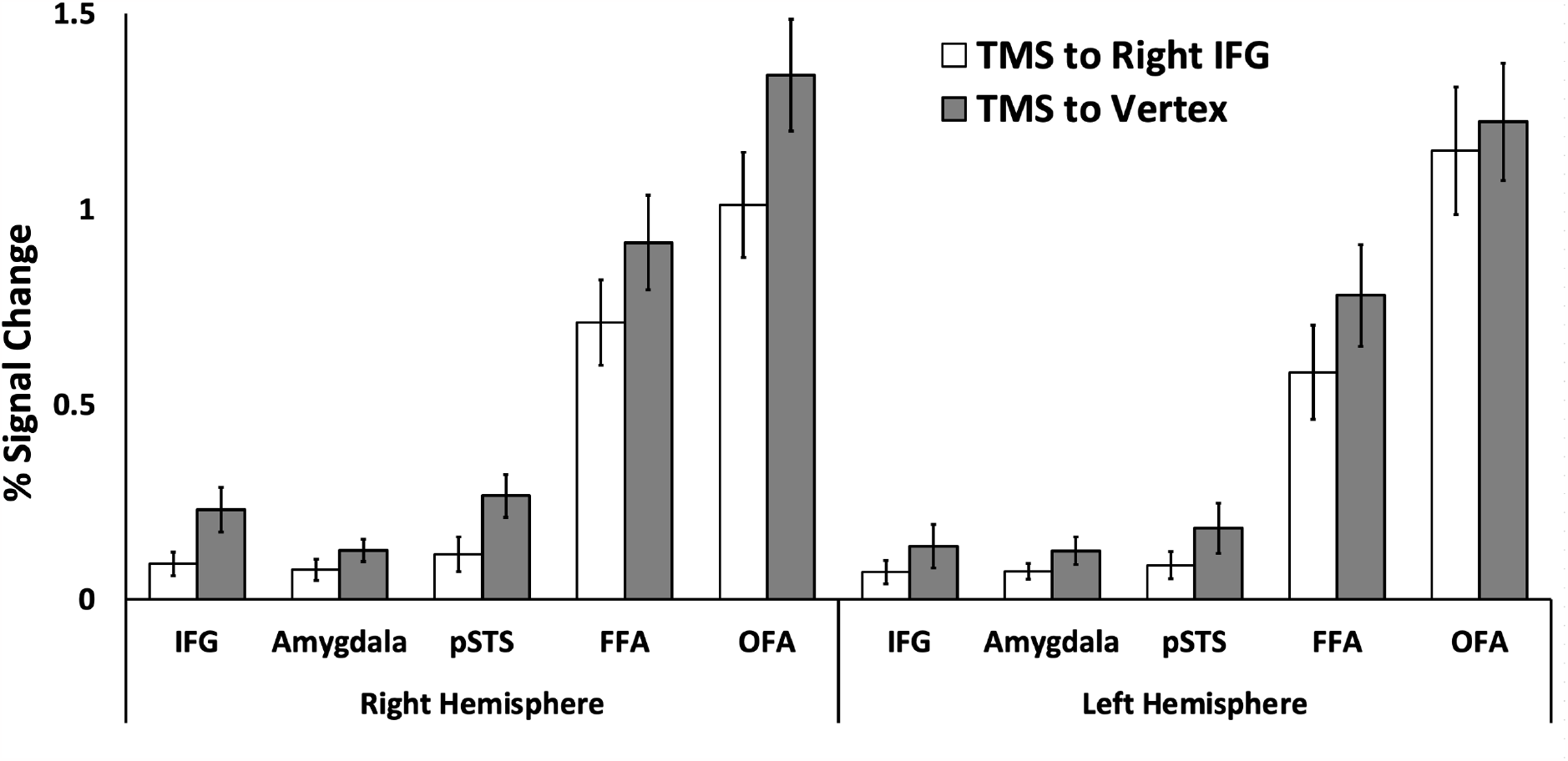
Results of the ROI analysis performed in face-selective areas for the facial expression naming task. Percent signal change (PSC) data for two stimulation conditions (right IFG and vertex) in the right and left IFG, amygdala, pSTS, FFA and OFA. Analyses revealed a significant main effect of stimulation (*p* = 0.015) in which TMS delivered over the right IFG reduced the neural response to expression naming across all nodes of the face network. There were no significant interactions. Error bars show standard errors of the mean across participants.

Data were entered into a 2 (stimulation: Right IFG, Vertex) by 2 (Hemisphere: right, left) by 5 (ROI: IFG, FFA, pSTS, OFA) repeated measures analysis of variance (ANOVA). Results showed significant main effects of stimulation (F (1,19) = 7.2, *p* = 0.015; partial η ^2^ = 0.275) and ROI (F (4,76) = 66.9, *p* < 0.001; partial η 2 = 0.779) but not of hemisphere (F (1,19) = 1.2, *p* = 0.27; partial η ^2^ = 0.063). There was no significant 3-way interaction between stimulation, hemisphere, and ROI (F (4,76) = 1.9, *p* = 0.112; partial η ^2^ = 0.09). All three of the two-way interactions were also non-significant (*p >* 0.065).

### Multivoxel pattern wide analysis (MVPA) Analysis

Finally, we performed a multivoxel pattern analysis (MVPA) on the facial expression data. This was done to establish whether TMS delivered over the right IFG selectively impaired the neural discriminability of the facial expressions presented (anger, disgust fear, happy, neutral, sad, surprise). Emotions could be discriminated from activity patterns in the bilateral pSTS, both after TMS over IFG (t (19) = 3.14, p = 0.003) and vertex (trending at t(19) = 1.59, p= 0.06). Emotions were also discriminable from activity patterns in the bilateral FFA (trending at t (19) = 1.37, p=0.09) and OFA (t (19) = 2.86, p = 0.005), but only after TMS over vertex. The OFA was the only region showing a TMS-related difference: Emotions were more readily discriminable from OFA response patterns when TMS was applied over vertex compared to TMS over the IFG (t (19) = 2.17, p = 0.04).

## Discussion

In the current study we disrupted the lateral PFC while participants were scanned using fMRI performing a facial expression naming task. Prior to scanning TMS was delivered over a functionally localized face-selective area centered at the right IFG, or over the vertex control site. The results of a whole brain group analysis (Figure 2) demonstrated that TMS delivered over the right IFG decreased neural activity in the left inferior frontal gyrus (compared to the vertex). The same analysis also revealed an increase in neural activity in three nodes of the default mode network: the right superior frontal gyrus (SFG), right angular gyrus and the bilateral middle cingulate gyrus. Our results are consistent with theories proposing that emotions are processed across functional brain networks that are specialized for different types of cognitive functions (Pessoa, 2018; Smallwood et al., 2021). The lateral PFC is a node in the central executive network necessary for cognitive control (Uddin, 2015). The DMN processes inner states such as mind wandering and inner thoughts (Smallwood et al., 2021). Our results demonstrate the dynamic nature of the interactions between these networks. Specifically, disrupting the right PFC with TMS increased the neural activity in the DMN while participants named facial expressions. This is consistent with a recent theory proposing a push / pull interaction between the central executive network and the DMN for emotion processing (Satpute & Lindquist, 2019).

The region of interest (ROI) analysis of the face-selective areas revealed a main effect of stimulation. TMS delivered over the right IFG reduced the neural response across all bilateral face ROIs compared to the vertex control condition (Figure 3). Our prior studies that combined TMS and fMRI also demonstrated distributed disruption across the face network (Groen et al., 2021; Handwerker et al., 2020; Pitcher et al., 2014; Pitcher et al., 2017). The lack of an interaction (that would have shown a greater impairment in selected face-selective areas than others) suggests that all five ROIs in both hemispheres are causally connected to the IFG during facial expression naming. This is consistent with patient and TMS studies showing that the IFG (Jastorff et al., 2016), pSTS (Pitcher, 2014; Sliwinska, Elson, et al., 2020), FFA (Rezlescu et al., 2012), OFA (Pitcher et al., 2008) and the amygdala (Adolphs et al., 1994) are all causally involved in facial expression recognition. Our findings reveal that IFG is directly or indirectly connected to all other regions in the face network.

The results of the group whole-brain analysis revealed that TMS delivered over the right IFG (compared the vertex control site) reduced neural activity in the left IFG for the expression naming task. The right IFG was selected as the TMS stimulation site because our prior study had demonstrated a greater response to visually presented faces in the right, more than the left IFG (Nikel et al., 2022). Despite this lateralisation the left frontal cortex has still been implicated in a range of face processing tasks. For example, prior fMRI studies have demonstrated that the left IFG exhibits greater activity in facial expression recognition tasks (Gorno-Tempini et al., 2001; Regenbogen et al., 2012; Trautmann et al., 2009). In addition, other tasks such as evaluating the social impact of facial expressions (Prochnow et al., 2014) and facial expression matching tasks (Sreenivas et al., 2012) also generate greater activity in the left IFG. The reduction in neural activity in the current study may have also been partially driven by the silent naming task participants performed. This would be consistent with the established role of the left IFG as a “high-level” language brain area (Fedorenko & Blank, 2020; Fedorenko & Thompson-Schill, 2014). More generally, our data show that face networks in both hemispheres are tightly interconnected, where disruption of one network node (the right IFG) has causal consequences on the activity in contralateral nodes like the left IFG.

We also performed an exploratory MVPA analysis to establish whether TMS disruption of the right IFG disrupted the neural representation of emotions. This analysis revealed that TMS over the IFG reduced the neural discriminability of emotions in OFA, but not any of the other regions. This suggests that TMS to the IFG can disrupt emotion processing in areas of the core face network. It is worth noting that this result was obtained under experimental conditions that are suboptimal for MVPA: The temporally constrained nature of TBS effects (lasting for only about 30min; Groen et al., 2021; Handwerker et al., 2020; Pitcher et al., 2014; Pitcher et al., 2017) drastically reduces the amount of available fMRI data, compared to typical MVPA studies of emotion processing (Harry et al., 2013; Said et al., 2010; Wegrzyn et al., 2015). Whether the effect observed in the OFA here truly extends to a larger set of areas, perhaps including the FFA, could be tested in future studies that use concurrent fMRI/TMS approaches (Mizutani-Tiebel et al., 2022) to increase the amount of available data. A surprising result in our MVPA is that pSTS, which is often considered a key region for emotion discrimination, did not show altered emotion representations after TMS to the IFG. Future studies should investigate whether such effects appear when emotion processing is probed with dynamic stimuli, which are strongly preferred by the region (Pitcher & Ungerleider, 2021).

The overall pattern of our results demonstrates that naming facial expressions is causally dependent on the interaction of different functional brain networks. While it is common for researchers to talk about the face processing network (Haxby et al., 2000) it is also important to note that the nodes of this network are distributed across brain areas with different cognitive functions. These include visual areas in occipito-temporal cortex (FFA, OFA, pSTS), emotion processing areas (the amygdala) and cognitive control areas (IFG). The results of the current study demonstrate the push / pull dynamic network interactions between these brain areas and nodes in the DMN. This is consistent with models of that proposing that emotion processing is a complex process that is dependent on the interactions of brain networks with different cognitive functions (Pessoa, 2018; Satpute & Lindquist, 2019; Uddin, 2015).

## References

Adolphs, R. (2002). Recognizing emotion from facial expressions: psychological and neurological mechanisms [Review]. Behavioral and cognitive neuroscience reviews, 1(1), 21–62. https://doi.org/10.1177/1534582302001001003

Adolphs, R., Tranel, D., Damasio, H., & Damasio, A. (1994). Impaired recognition of emotion in facial expressions following bilateral damage to the human amygdala [Article]. Nature, 372(6507), 669–672. https://doi.org/10.1038/372669a0

Baldauf, D., & Desimone, R. (2014). Neural mechanisms of object-based attention. Science, 344(6182), 424–427. https://doi.org/10.1126/science.1247003

Barrett, L. F., & Satpute, A. B. (2013). Large-scale brain networks in affective and social neuroscience: towards an integrative functional architecture of the brain. Curr Opin Neurobiol, 23(3), 361–372. https://doi.org/10.1016/j.conb.2012.12.012

Calder, A. J., & Young, A. W. (2005). Understanding the recognition of facial identity and facial expression [Review]. Nature Reviews Neuroscience, 6(8), 641–651. https://doi.org/10.1038/nrn1724

Chan, A. W. (2013). Functional organization and visual representations of human ventral lateral prefrontal cortex. Front Psychol, 4, 371. https://doi.org/10.3389/fpsyg.2013.00371

Courtney, S. M., Ungerleider, L. G., Keil, K., & Haxby, J. V. (1996). Object and spatial visual working memory activate separate neural systems in human cortex. Cereb Cortex, 6(1), 39–49. https://doi.org/10.1093/cercor/6.1.39

Davies-Thompson, J., & Andrews, T. J. (2012). Intra- and interhemispheric connectivity between face-selective regions in the human brain. J Neurophysiol, 108(11), 3087–3095. https://doi.org/10.1152/jn.01171.2011

Eickhoff, S. B., Stephan, K. E., Mohlberg, H., Grefkes, C., Fink, G. R., Amunts, K., & Zilles, K. (2005). A new SPM toolbox for combining probabilistic cytoarchitectonic maps and functional imaging data. NeuroImage, 25(4), 1325–1335. https://doi.org/10.1016/j.neuroimage.2004.12.034

Fedorenko, E., & Blank, I. A. (2020). Broca’s Area Is Not a Natural Kind. Trends Cogn Sci, 24(4), 270–284. https://doi.org/10.1016/j.tics.2020.01.001

Fedorenko, E., & Thompson-Schill, S. L. (2014). Reworking the language network. Trends Cogn Sci, 18(3), 120–126. https://doi.org/10.1016/j.tics.2013.12.006

Gauthier, I., Tarr, M. J., Moylan, J., Skudlarski, P., Gore, J. C., & Anderson, A. W. (2000). The fusiform ‘face area’ is part of a network that processes faces at the individual level [Article]. Journal of Cognitive Neuroscience, 12(3), 495–504. https://doi.org/10.1162/089892900562165

Goldsworthy, M. R., Pitcher, J. B., & Ridding, M. C. (2012). A comparison of two different continuous theta burst stimulation paradigms applied to the human primary motor cortex [Article]. Clinical Neurophysiology, 123(11), 2256–2263. https://doi.org/10.1016/j.clinph.2012.05.001

Gorno-Tempini, M. L., Pradelli, S., Serafini, M., Pagnoni, G., Baraldi, P., Porro, C., Nicoletti, R., Umita, C., & Nichelli, P. (2001). Explicit and incidental facial expression processing: an fMRI study. NeuroImage, 14(2), 465–473. https://doi.org/10.1006/nimg.2001.0811

Groen, I. I. A., Silson, E. H., Pitcher, D., & Baker, C. I. (2021). Theta-burst TMS of lateral occipital cortex reduces BOLD responses across category-selective areas in ventral temporal cortex. NeuroImage, 230, 117790. https://doi.org/10.1016/j.neuroimage.2021.117790

Handwerker, D. A., Ianni, G., Gutierrez, B., Roopchansingh, V., Gonzalez-Castillo, J., Chen, G., Bandettini, P. A., Ungerleider, L. G., & Pitcher, D. (2020). Theta-burst TMS to the posterior superior temporal sulcus decreases resting-state fMRI connectivity across the face processing network. Netw Neurosci, 4(3), 746–760. https://doi.org/10.1162/netn_a_00145

Harry, B., Williams, M. A., Davis, C., & Kim, J. (2013). Emotional expressions evoke a differential response in the fusiform face area. Front Hum Neurosci, 7, 692. https://doi.org/10.3389/fnhum.2013.00692

Haxby, J. V., Gobbini, M. I., Furey, M. L., Ishai, A., Schouten, J. L., & Pietrini, P. (2001). Distributed and overlapping representations of faces and objects in ventral temporal cortex [Article]. Science, 293(5539), 2425–2430. https://doi.org/10.1126/science.1063736

Haxby, J. V., Hoffman, E. A., & Gobbini, M. I. (2000). The distributed human neural system for face perception [Review]. Trends in Cognitive Sciences, 4(6), 223–233. https://doi.org/10.1016/S1364-6613(00)01482-0

Heekeren, H. R., Marrett, S., Bandettini, P. A., & Ungerleider, L. G. (2004). A general mechanism for perceptual decision-making in the human brain. Nature, 431(7010), 859–862. https://doi.org/10.1038/nature02966

Huang, Y. Z., Edwards, M. J., Rounis, E., Bhatia, K. P., & Rothwell, J. C. (2005). Theta burst stimulation of the human motor cortex [Article]. Neuron, 45(2), 201–206. https://doi.org/10.1016/j.neuron.2004.12.033

Iidaka, T., Omori, M., Murata, T., Kosaka, H., Yonekura, Y., Okada, T., & Sadato, N. (2001). Neural interaction of the amygdala with the prefrontal and temporal cortices in the processing of facial expressions as revealed by fMRI. J Cogn Neurosci, 13(8), 1035–1047. https://doi.org/10.1162/089892901753294338

Ishai, A., Haxby, J. V., & Ungerleider, L. G. (2002). Visual imagery of famous faces: Effects of memory and attention revealed by fMRI [Article]. NeuroImage, 17(4), 1729–1741. https://doi.org/10.1006/nimg.2002.1330

Jastorff, J., De Winter, F. L., Van den Stock, J., Vandenberghe, R., Giese, M. A., & Vandenbulcke, M. (2016). Functional dissociation between anterior temporal lobe and inferior frontal gyrus in the processing of dynamic body expressions: Insights from behavioral variant frontotemporal dementia. Human Brain Mapping, 37(12), 4472–4486. https://doi.org/10.1002/hbm.23322

Kanwisher, N., McDermott, J., & Chun, M. M. (1997). The fusiform face area: A module in human extrastriate cortex specialized for face perception [Article]. Journal of Neuroscience, 17(11), 4302–4311. https://doi.org/10.1523/jneurosci.17-11-04302.1997

Küçük, E., Foxwell, M., Kaiser, D., & Pitcher, D. (2022). Moving and static faces, bodies, objects and scenes are differentially represented across the three visual pathways. bioRxiv, 2022.2011.2030.518408. https://doi.org/10.1101/2022.11.30.518408

Lanzoni, L., Ravasio, D., Thompson, H., Vatansever, D., Margulies, D., Smallwood, J., & Jefferies, E. (2020). The role of default mode network in semantic cue integration. NeuroImage, 219, 117019. https://doi.org/10.1016/j.neuroimage.2020.117019

Lindquist, K. A., Wager, T. D., Kober, H., Bliss-Moreau, E., & Barrett, L. F. (2012). The brain basis of emotion: a meta-analytic review. Behav Brain Sci, 35(3), 121–143. https://doi.org/10.1017/S0140525X11000446

McCarthy, G., Puce, A., Gore, J. C., & Allison, T. (1997). Face-specific processing in the human fusiform gyrus [Article]. Journal of Cognitive Neuroscience, 9(5), 605–610. https://doi.org/10.1162/jocn.1997.9.5.605

Mizutani-Tiebel, Y., Tik, M., Chang, K. Y., Padberg, F., Soldini, A., Wilkinson, Z., Voon, C. C., Bulubas, L., Windischberger, C., & Keeser, D. (2022). Concurrent TMS-fMRI: Technical Challenges, Developments, and Overview of Previous Studies. Front Psychiatry, 13, 825205. https://doi.org/10.3389/fpsyt.2022.825205

Nikel, L., Sliwinska, M. W., Kucuk, E., Ungerleider, L. G., & Pitcher, D. (2022). Measuring the response to visually presented faces in the human lateral prefrontal cortex. Cereb Cortex Commun, 3(3), tgac036. https://doi.org/10.1093/texcom/tgac036

Nyffeler, T., Wurtz, P., Luscher, H. R., Hess, C. W., Senn, W., Pflugshaupt, T., von Wartburg, R., Luthi, M., & Muri, R. M. (2006). Repetitive TMS over the human oculomotor cortex: comparison of 1-Hz and theta burst stimulation. Neurosci Lett, 409(1), 57–60. https://doi.org/10.1016/j.neulet.2006.09.011

Oosterhof, N. N., Connolly, A. C., & Haxby, J. V. (2016). CoSMoMVPA: Multi-Modal Multivariate Pattern Analysis of Neuroimaging Data in Matlab/GNU Octave. Front Neuroinform, 10, 27. https://doi.org/10.3389/fninf.2016.00027

Pessoa, L. (2018). Understanding emotion with brain networks. Curr Opin Behav Sci, 19, 19–25. https://doi.org/10.1016/j.cobeha.2017.09.005

Pitcher, D. (2014). Facial expression recognition takes longer in the posterior superior temporal sulcus than in the occipital face area [Article]. Journal of Neuroscience, 34(27), 9173–9177. https://doi.org/10.1523/JNEUROSCI.5038-13.2014

Pitcher, D., Dilks, D. D., Saxe, R. R., Triantafyllou, C., & Kanwisher, N. (2011). Differential selectivity for dynamic versus static information in face-selective cortical regions [Article]. NeuroImage, 56(4), 2356–2363. https://doi.org/10.1016/j.neuroimage.2011.03.067

Pitcher, D., Duchaine, B., & Walsh, V. (2014). Combined TMS and fMRI reveal dissociable cortical pathways for dynamic and static face perception [Article]. Current Biology, 24(17), 2066–2070. https://doi.org/10.1016/j.cub.2014.07.060

Pitcher, D., Garrido, L., Walsh, V., & Duchaine, B. C. (2008). Transcranial magnetic stimulation disrupts the perception and embodiment of facial expressions [Article]. Journal of Neuroscience, 28(36), 8929–8933. https://doi.org/10.1523/JNEUROSCI.1450-08.2008

Pitcher, D., Japee, S., Rauth, L., & Ungerleider, L. G. (2017). The superior temporal sulcus is causally connected to the amygdala: A combined TBS-fMRI study [Article]. Journal of Neuroscience, 37(5), 1156–1161. https://doi.org/10.1523/JNEUROSCI.0114-16.2016

Pitcher, D., & Ungerleider, L. G. (2021). Evidence for a Third Visual Pathway Specialized for Social Perception [Review]. Trends in Cognitive Sciences, 25(2), 100–110. https://doi.org/10.1016/j.tics.2020.11.006

Prochnow, D., Brunheim, S., Steinhauser, L., & Seitz, R. J. (2014). Reasoning about the implications of facial expressions: A behavioral and fMRI study on low and high social impact. Brain and Cognition, 90, 165–173. https://doi.org/10.1016/j.bandc.2014.07.004

Puce, A., Allison, T., Asgari, M., Gore, J. C., & McCarthy, G. (1996). Differential sensitivity of human visual cortex to faces, letterstrings, and textures: A functional magnetic resonance imaging study [Article]. Journal of Neuroscience, 16(16), 5205–5215. https://doi.org/10.1523/jneurosci.16-16-05205.1996

Puce, A., Allison, T., Bentin, S., Gore, J. C., & McCarthy, G. (1998). Temporal cortex activation in humans viewing eye and mouth movements [Article]. Journal of Neuroscience, 18(6), 2188–2199. https://doi.org/10.1523/jneurosci.18-06-02188.1998

Raichle, M. E. (2015). The Brain’s Default Mode Network. In Annual Review of Neuroscience (Vol. 38, pp. 433–447): Annual Reviews Inc.

Regenbogen, C., Schneider, D. A., Gur, R. E., Schneider, F., Habel, U., & Kellermann, T. (2012). Multimodal human communication--targeting facial expressions, speech content and prosody. NeuroImage, 60(4), 2346–2356. https://doi.org/10.1016/j.neuroimage.2012.02.043

Renzi, C., Schiavi, S., Carbon, C. C., Vecchi, T., Silvanto, J., & Cattaneo, Z. (2013). Processing of featural and configural aspects of faces is lateralized in dorsolateral prefrontal cortex: a TMS study. NeuroImage, 74, 45–51. https://doi.org/10.1016/j.neuroimage.2013.02.015

Rezlescu, C., Pitcher, D., & Duchaine, B. (2012). Acquired prosopagnosia with spared within-class object recognition but impaired recognition of degraded basic-level objects [Article]. Cognitive Neuropsychology, 29(4), 325–347. https://doi.org/10.1080/02643294.2012.749223

Said, C. P., Moore, C. D., Engell, A. D., Todorov, A., & Haxby, J. V. (2010). Distributed representations of dynamic facial expressions in the superior temporal sulcus [Article]. Journal of Vision, 10(5), Article 11. https://doi.org/10.1167/10.5.11

Satpute, A. B., & Lindquist, K. A. (2019). The Default Mode Network’s Role in Discrete Emotion. Trends in Cognitive Sciences, 23(10), 851–864. https://doi.org/10.1016/j.tics.2019.07.003

Seeley, W. W., Menon, V., Schatzberg, A. F., Keller, J., Glover, G. H., Kenna, H., Reiss, A. L., & Greicius, M. D. (2007). Dissociable intrinsic connectivity networks for salience processing and executive control. J Neurosci, 27(9), 2349–2356. https://doi.org/10.1523/JNEUROSCI.5587-06.2007

Sliwinska, M. W., Bearpark, C., Corkhill, J., McPhillips, A., & Pitcher, D. (2020). Dissociable pathways for moving and static face perception begin in early visual cortex: Evidence from an acquired prosopagnosic [Article]. Cortex, 130, 327–339. https://doi.org/10.1016/j.cortex.2020.03.033

Sliwinska, M. W., Elson, R., & Pitcher, D. (2020). Dual-site TMS demonstrates causal functional connectivity between the left and right posterior temporal sulci during facial expression recognition [Article]. Brain Stimulation, 13(4), 1008–1013. https://doi.org/10.1016/j.brs.2020.04.011

Sliwinska, M. W., Searle, L. R., Earl, M., O’Gorman, D., Pollicina, G., Burton, A. M., & Pitcher, D. (2022). Face learning via brief real-world social interactions includes changes in face-selective brain areas and hippocampus. Perception, 3010066221098728. https://doi.org/10.1177/03010066221098728

Smallwood, J., Bernhardt, B. C., Leech, R., Bzdok, D., Jefferies, E., & Margulies, D. S. (2021). The default mode network in cognition: a topographical perspective. Nat Rev Neurosci, 22(8), 503–513. https://doi.org/10.1038/s41583-021-00474-4

Sreenivas, S., Boehm, S. G., & Linden, D. E. (2012). Emotional faces and the default mode network. Neurosci Lett, 506(2), 229–234. https://doi.org/10.1016/j.neulet.2011.11.012

Trautmann, S. A., Fehr, T., & Herrmann, M. (2009). Emotions in motion: Dynamic compared to static facial expressions of disgust and happiness reveal more widespread emotion-specific activations [Article]. Brain Research, 1284, 100–115. https://doi.org/10.1016/j.brainres.2009.05.075

Tsuchida, A., & Fellows, L. K. (2012). Are You Upset? Distinct Roles for Orbitofrontal and Lateral Prefrontal Cortex in Detecting and Distinguishing Facial Expressions of Emotion. Cerebral Cortex, 22(12), 2904–2912. https://doi.org/10.1093/cercor/bhr370

Uddin, L. Q. (2015). Salience processing and insular cortical function and dysfunction. Nat Rev Neurosci, 16(1), 55–61. https://doi.org/10.1038/nrn3857

Wager, T. D., Kang, J., Johnson, T. D., Nichols, T. E., Satpute, A. B., & Barrett, L. F. (2015). A Bayesian model of category-specific emotional brain responses. PLoS Comput Biol, 11(4), e1004066. https://doi.org/10.1371/journal.pcbi.1004066

Wang, Y., Metoki, A., Smith, D. V., Medaglia, J. D., Zang, Y., Benear, S., Popal, H., Lin, Y., & Olson, I. R. (2020). Multimodal mapping of the face connectome. Nat Hum Behav, 4(4), 397–411. https://doi.org/10.1038/s41562-019-0811-3

Wegrzyn, M., Riehle, M., Labudda, K., Woermann, F., Baumgartner, F., Pollmann, S., Bien, C. G., & Kissler, J. (2015). Investigating the brain basis of facial expression perception using multi-voxel pattern analysis. Cortex, 69, 131–140. https://doi.org/10.1016/j.cortex.2015.05.003

